# Serotypes of Dengue Virus and *Kdr* Mutations (*F1534C, S989P, V410L, V1016I,* and *V1016G*) in *Aedes aegypti* and *Aedes albopictus* Mosquitoes from the commune of Abomey-Calavi, Southern Benin

**DOI:** 10.1101/2025.06.20.660662

**Authors:** Tatchémè Filémon Tokponnon, Oloubo Djibril Amoussa, Zoulkifilou Sare Dabou, Nicodéme Kpemasse, Minassou Juvenal Ahouandjinou, Linda Towakinou, Brunelle Agassounon, Olivier Tandjekpon, Fabrice Gandaho, Mistourath Issa, Idayath Gounou Yérima, Festus Houessinon, Victorien Dougnon, Richard M. Oxborough, Louisa Mesenger, Razaki Osse, Lamine Baba-Moussa, Dorothée Kinde Gazard, Martin Akogbeto

## Abstract

Dengue is a viral infection transmitted by mosquitoes of the *Aedes* genus, responsible for millions of cases each year. In Benin, several outbreaks have been reported, particularly in Abomey-Calavi, where the first dengue-related death was recorded in 2019. In the absence of a specific treatment, vector control remains the primary preventive measure. However, the emergence of insecticide resistance, notably through *kdr* mutations, could compromise its effectiveness.

This study aims to identify the dengue virus serotypes circulating in the commune of Abomey-Calavi and to assess the frequency of *kdr* mutations *V410L, V1016G, V1016I, F1534C, and S989P* in *Aedes aegypti* and *Aedes albopictus* mosquitoes. Mosquitoes were collected in August 2024 from four arrondissement Togba, Calavi, Akassato, and Godomey through morning and afternoon spraying. After morphological identification, molecular analyses were conducted to detect the viral serotypes and *kdr* mutations.

A total of 218 *Aedes* mosquitoes were collected, with a predominance of *Aedes aegypti* (68.8%) compared to *Aedes albopictus* (31.2%). The overall dengue positivity rate was 23.39% (95% CI: 17.9 - 29.6). The most frequent serotype was DENV-3 (12.4%), followed by DENV-2 (6.9%), DENV-1 (5%), and DENV-4 (0.09%). Six mosquitoes were co-infected with two serotypes, including three with DENV-1 and DENV-4. *kdr* mutations were detected in both species, with frequencies of 45% for *V1016G,* 44% for *S989P,* 40% for *F1534C*, 22% for *V410L*, and 19% for *V1016I.* Four mosquitoes carried three simultaneous mutations, while twenty-two carried two. Two mosquitoes co-infected with two serotypes also carried two different mutations. These results highlight the active circulation of the dengue virus and the presence of *kdr* mutations in *Aedes* mosquitoes in Abomey-Calavi. However, no significant association was observed between dengue infection and *kdr* mutations, and their distribution was independent of the viral serotype. These findings emphasize the need for regular monitoring of *kdr* mutations to adapt vector control strategies and limit the spread of dengue.

## Introduction

Dengue is an acute viral infection caused by a single-stranded RNA virus classified in the *Flavivirus* genus. This infection poses a major public health challenge and significantly hinders socio-economic development, particularly in tropical and subtropical regions [1]. It is the most widespread mosquito-borne viral infection in the world and is primarily transmitted by *Aedes aegypti* and *Aedes albopictus* mosquitoes [2]. Each year, approximately 390 million people are infected with the dengue virus (DENV), of which 96 million develop symptoms [3,4]. Dengue generally presents with mild symptoms such as fever, but it can progress to severe forms requiring hospitalization [5]. Among those who develop symptoms, nearly 25,000 die annually worldwide [3]. The World Health Organization (WHO) has observed a significant increase in dengue cases in recent decades. Today, more than one-third of the global population lives in endemic areas where dengue poses a major public health threat [6]. In Africa, dengue has been detected in 34 countries, and West Africa is considered a high-risk area due to sanitary conditions favorable to the proliferation of *Aedes* mosquitoes [7]. In West Africa, all four dengue virus serotypes (DENV1, DENV2, DENV3, and DENV4) are actively circulating [8]. Nigeria and Burkina Faso, both neighboring countries of Benin, are recognized as endemic areas [9,10]. The first officially reported case of dengue in Benin occurred in 2019 [11]. Today, several studies in Benin have confirmed the presence of *Aedes aegypti*, the primary dengue vector. For instance, studies by Yadouleton *et al.* (2018) and Padonou *et al.* (2023) showed a high predominance of *Aedes aegypti* in Cotonou and Avrankou, respectively [12,13]. The first suspected dengue cases in Benin were reported in 1987 among German humanitarian workers [14]. Since then, different serotypes have been identified in the country, in both humans and mosquitoes [11,15]. More recently, in 2022, the presence of serotype 3 was confirmed in mosquitoes in Benin [16]. These observations indicate active, and likely underestimated, circulation of several serotypes in Benin.

To date, there is neither a specific treatment nor an effective vaccine against dengue, and vector control remains the only method for reducing morbidity and mortality linked to the disease [17]. In Asia and South America, surveillance and control of *Aedes* mosquitoes are integral parts of arbovirus prevention strategies. In contrast, in Africa, vector control is generally reactive and limited to interventions during outbreaks [18]. Vector control programs in Africa mainly focus on *Anopheles* vectors and rely on indoor residual spraying of insecticides and insecticide-treated bed nets [19]. Among the major classes of insecticides used in public health, pyrethroids are the most widely used [20]. However, their excessive use in vector control and agriculture has led to insecticide resistance in *Anopheles* species [21]. These effects are less well documented in *Aedes* mosquitoes, whose ecology and behavior differ. Knowledge of insecticide resistance and the underlying mechanisms in *Aedes* mosquitoes remains limited in Africa [21].

Insecticide resistance mechanisms are complex. Among them, mutations in the voltage-gated sodium channel (VGSC) gene in *Aedes aegypti* are the most studied. Numerous knockdown resistance (*Kdr*) mutations in the VGSC gene have been identified worldwide, including *V1016I, F174I, E478K, S723T, D1763Y, Q1853R, V410L, S989P, I1011V, V1016G, I1011M*, and *F1534C* [22]. Among these, three specific mutations *S989P, V1016G*, and *F1534C* have been reported in Benin [23]. The resurgence of dengue outbreaks highlights the need for effective vector control programs. However, to our knowledge, no study in Benin has simultaneously investigated the presence of different dengue serotypes and the *Kdr* mutations in *Aedes aegypti* and *Aedes albopictus* populations. Furthermore, WHO recommends studies that integrate both the transmission level of vector-borne diseases and the resistance status of their vectors [24]. In this context, our research focused on the detection of dengue virus serotypes and *Kdr* mutations *V410L, V1016G, V1016I, S989P*, and *F1534C* associated with insecticide resistance in *Aedes aegypti* and *Aedes albopictus* mosquitoes in the commune of Abomey-Calavi. The results of this study will help assess the dengue situation in the area and guide vector control strategies while contributing to the epidemiological surveillance of the disease.

## Materials and Methods

### Sampling Site

The study was conducted in the **c**ommune of Abomey-Calavi, in southern Benin, specifically in the arrondissement of Calavi, Godomey, Akassato, and Togba (Figure 1), due to its high population density and previously reported dengue cases [11]. The minimum sample size of households was calculated using Schwartz’s formula, based on an estimated prevalence of 87.7% from a study by Padonou et al. (2024), resulting in a required total of 166 households [25]. However, 252 households were surveyed, with variable distribution across the arrondissement. The higher participation in Calavi was due to better awareness among residents, most of whom were students. Mosquito collection took place in August to September 2024, during the short dry season. An indoor spraying method using the insecticide “Boxer” was employed early in the morning and in the evening. Participating households were randomly selected, and informed and consenting residents responded to a questionnaire. Prior to spraying, rooms were closed and covered with white sheets to facilitate the collection of fallen mosquitoes. After the application of the insecticide, moribund or dead mosquitoes were collected and transported to the laboratory for analysis.

**Figure 1.**
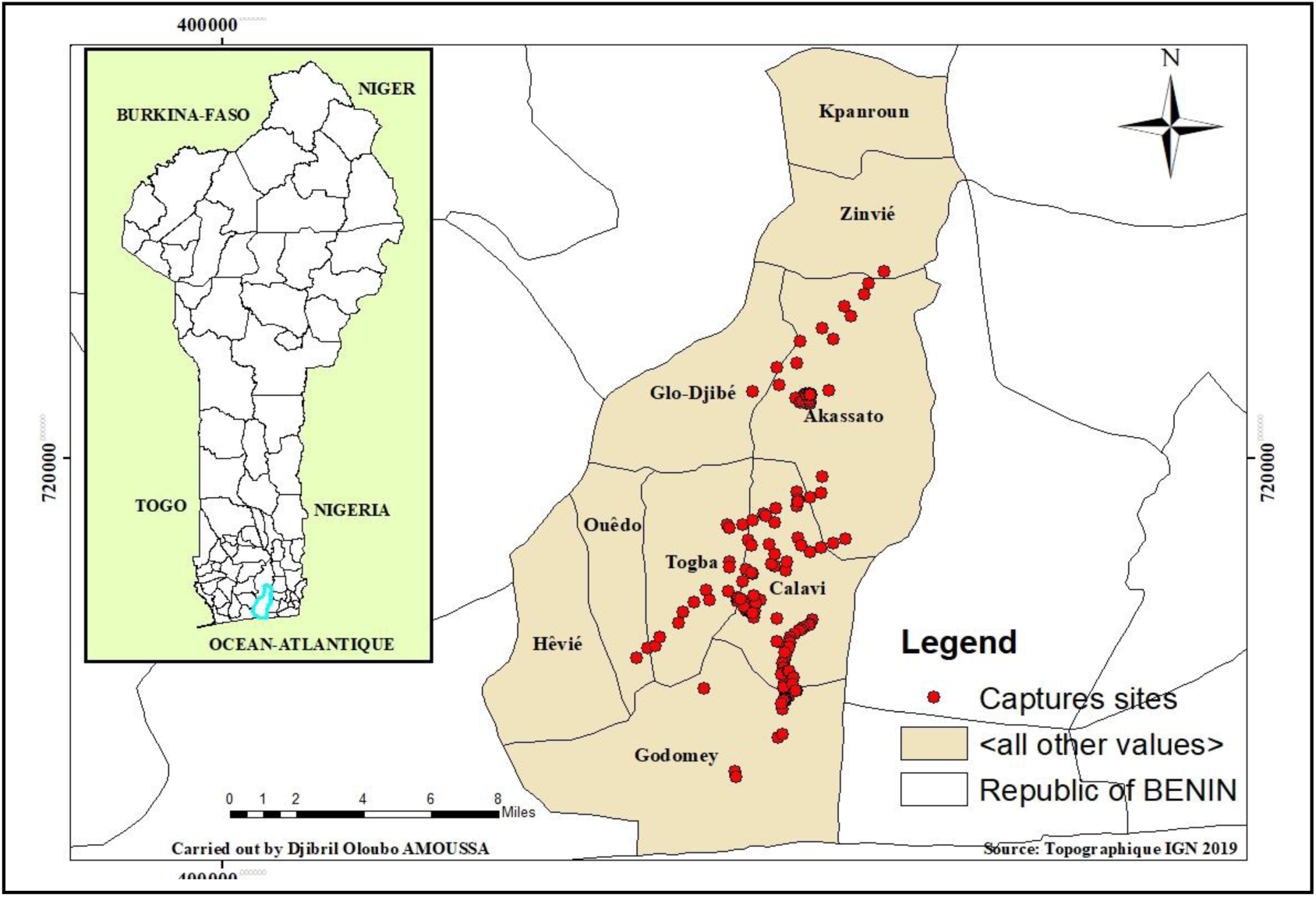
Map of spraying sites Abomey-Calavi, Benin. The dots correspond to different households in Abomey-Calavi

### 2.2. Morphological Identification and Processing of *Aedes* Mosquitoes

Collected mosquitoes were morphologically identified using the taxonomic keys of Fontenille and collaborators [26]. *Aedes albopictus* mosquitoes were distinguished by a central line of white scales on the thorax, while *Aedes aegypti* displayed a lyre-shaped pattern. Specimens were preserved at -80°C in RNA later for further analysis. Each mosquito was dissected into two parts: the head, thorax, and abdomen were stored in RNA later for dengue virus and serotype detection, while wings and legs were used for *Kdr* mutation analysis.

### 2.3. RNA Extraction and cDNA Synthesis

RNA was extracted using the Qiagen RNeasy 96 Kit® (Qiagen, UK). The head, thorax, and abdomen of each mosquito were ground in 150 µl of RTL buffer using the Qiagen Tissue Lyser II. Then, 150 µl of 70% ethanol were added to each tube, and the supernatant was transferred onto the RNeasy 96 column plate. The plates were sealed with an AirPore tape sheet and centrifuged at 6000 rpm (∼5600 x g) for 2 minutes. A three-step washing process followed: 100 µL of RW1 buffer with 2-minute centrifugation, then two washes with 100 µL of RPE buffer, with 2-minute and 4-minute spins respectively. After washing, the RNeasy 96 plates were placed onto 96-well elution plates. Then, 45 µL of RNase-free water were added to each well, incubated at room temperature for 1 minute, and centrifuged again at 5600 g for 2 minutes. RNA concentration was measured with a Nanodrop, and samples were stored at - 80°C.

### 2.4. DNase Treatment and Complementary DNA Synthesis

Dengue virus and serotype detection followed the nested PCR protocol described by Pérez and collaborators [27]. This method involves two amplification steps: the first confirms the presence of the virus, and the second identifies the serotype. The first PCR was performed using an Eppendorf Nexus thermocycler and the HotStarTaq Mastermix kit (Qiagen). The 25 µL reaction mixture included 2 µL of cDNA, 2 µL of each primer (mD1 and D2), 2.5 µL of 10X buffer, 0.5 µL of MgCl₂, 14.4 µL of distilled water, 1 µL of Q buffer, and 0.1 µL of HotStarTaq polymerase. The reaction started with denaturation at 95°C for 15 minutes, followed by 35 cycles of 15 seconds at 94°C (denaturation), 15 seconds at 55°C (annealing), and 30 seconds at 72°C (extension), ending with a final extension at 72°C for 10 minutes. The first step was expected to produce a DNA fragment of 511 base pairs.

The second PCR to identify dengue virus serotypes used the same HotStarTaq mastermix. The 25 µL mix included 5 µL of the first-round PCR product, 2 µL of each specific primer (mD1, rTS1, mTS2, TS3, and rTS4), 2.5 µL of 10X buffer, 0.5 µL of MgCl₂, 1 µL of dNTPs, 0.1 µL of Taq polymerase, and 5.9 µL of distilled water. The program began with activation at 95°C for 15 minutes, followed by 25 cycles of 15 seconds at 95°C, 15 seconds at 55°C, and 30 seconds at 72°C, ending with a final extension at 72°C for 7 minutes. The PCR products were separated using 2% agarose gel electrophoresis and visualised with ethidium bromide staining. Serotypes were identified based on fragment sizes: 208 bp for DENV-1, 119 bp for DENV-2, 288 bp for DENV-3, and 260 bp for DENV-4. Primer sequences used for amplification were:

- mD1: 5’-TCAATATGCTGAAACGCGAGAAACCG-3’
- D2: 5’-TTGCACCAACAGTCAATGTCTTCAGGTTC-3’
- rTS1: 5’-CCCGTAACACTTTGATCGCT-3’
- mTS2: 5’-CGCACAAGGGCATGAACAGTTT-3’
- TS3: 5’-TAACATCATCATGAGACAGAGC-3’
- rTS4: 5’-TTCTCCCGTTCAGGATGTC-3’

### 2.5. Dengue Epidemic Risk Assessment in the Study Area

The House Index (HI) was used to assess dengue circulation risk in the human population [28]. This index corresponds to the percentage of positive houses out of all inspected houses. According to interpretation thresholds, an HI below 4% indicates low epidemiological risk, while an HI above 35% suggests high arbovirus transmission risk. An HI between 4% and 35% is considered moderate.

### 2.6. DNA Extraction and Determination of *Kdr* Mutations *F1534C, S989P, V410L, V1016I, and V1016G*

Genomic DNA was extracted using the method described by Tokponnon and collaborators [23]. The wings and legs of each specimen were ground with beads in 200 µL of 2% CTAB, then incubated at 65 °C for five minutes. Then, 200 µL of chloroform were added, after which the sample was centrifuged at 12,000 rpm. The resulting supernatant was collected and precipitated with 200 µL of isopropanol. Following a further centrifugation step, the DNA was washed with 70% ethanol, dried and resuspended in 40 µL of sterile water.

### 2.7. Allele-Specific PCR (AS-PCR) Genotyping

*Kdr* mutations *S989P, V1016G, F1534C* were detected using the protocol by Li and collaborators [29]. Each sample was tested with two primers: one targeting the wild-type allele and the other targeting the mutation. The primers used were:

- *S989P* (F: 5′AATGATATTAACAAAATTGCGC3′, R:5′GCACGCCTCTAATATTGATGC)
- *V1016G* (F: 5′GCCACCGTAGTGATAGGAAATC3′, R: 5′CGGGTTAAGTTTCGTTTAGTAGC3′)
- *F1534C* (F: 5′GGAGAACTACACGTGGGAGAAC3′, R: 5′CGCCACTGAAATTGAGAATAGC3′)

PCR was conducted in a 25 µL final volume with 2.5 µL of 10X buffer, 1.5 µL of MgCl₂ (1.5 mM), 1.5 µL of dNTPs (0.3 mM), 0.2 µL of Taq polymerase, and 9.8 µL of water and DNA. Each primer (forward, reverse, sensitive/mutant) was added at 2.5 µL. PCR conditions included initial denaturation at 94°C for 3 minutes, followed by 35 cycles of 30 seconds at 94°C (denaturation), annealing at 60°C (*F1534C* and *V1016G*) or 62°C (*S989P*), and 1 minute at 72°C (extension), with a final extension at 72°C for 7 minutes.

### 2.8. Detection of V410L and V1016I Mutations by Multiplex PCR

Multiplex PCR was used to detect wild-type and mutant alleles of *V410L* and *V1016I* mutations simultaneously, following the method by Kosinski and collaborators [22]. The 25 µL reaction mix included 13.8 µL of water, 2.5 µL of 10X buffer, 1.5 µL of dNTPs, 0.2 µL of Taq polymerase, and 2 µL of DNA. Each primer (forward and reverse) was added at 2.5 µL.

PCR included an initial denaturation at 94°C for 3 minutes, followed by 45 cycles of 30 seconds at 94°C, 30 seconds annealing at 56°C, and 60 seconds of extension at 72°C, ending with a final extension at 72°C for 10 minutes. The amplified fragment sizes were: 113 bp (*V410L* resistant), 133 bp (*V410L* sensitive), 82 bp (*V1016I* resistant), and 102 bp (*V1016I* sensitive).

### 2.9. Migration of PCR Products

Electrophoresis of PCR products was performed on a 2% agarose gel. The gel was prepared by dissolving 4 g of agarose in 200 mL of buffer, heating it in a microwave, and then adding 10 µL of BET. After cooling and casting the gel, wells were formed using a comb. PCR products were mixed with bromophenol blue and loaded into the wells (10 µL per sample). A molecular weight marker (100 bp) was loaded into the outermost wells. After electrophoresis, the fragments were visualized under UV light to check their size.

### 2.10. Statistical Analyses

The GPS coordinates of the households included in the study were recorded using the Kobocollect Android app. All analyses were performed using SPSS 21 and Excel 2019.

For *Kdr* mutation data, the allelic frequency of a biallelic gene (R and S) was estimated using the formula Fr(R) = (2nRR + nRS)/(2(nRR + nRS + nSS)), where Fr represents the frequency and n the number. This method was used to assess the frequency of mutations in the studied genes. Fisher’s exact test and chi-squared test were used to calculate p-values and identify significant differences.

### 2.11. Ethics approval and consent to participate

The protocol for this study was reviewed and approved by the CREC Institutional Research Ethics Committee (N°05/CEICREC/SA, Approval of 01/06/2024). Consent to participate in the study was obtained from the participants after they were fully informed about the study objectives, procedures, and any potential risks. Verbal consent was obtained, following a detailed explanation of the entire study protocol, as approved by the Ethics Committee.

## 3. Results

### 3.1. Diversity of Mosquitoes Captured in the Study Area

The study included 252 households, allowing the capture of 1,213 mosquitoes belonging to the genera *Mansonia*, *Culex*, *Aedes*, and *Anopheles*. The results show that *Mansonia africana* was the most represented species (53.66%), followed by *Aedes aegypti* (12.36%), *Culex annulioris* (9.23%), and *Aedes albopictus* (5.60%). The least frequent species was *Anopheles phaorensis* (0.16%).

A specific analysis of *Aedes aegypti* and *Aedes albopictus*, known dengue vectors, revealed that most specimens were collected in the arrondissement of Calavi (36.69%), followed by Godomey (25.68%), Akassato (23.39%), and Togba (14.22%). In all arrondissement, *Aedes aegypti* was more abundant than *Aedes albopictus* (table I).

**Table I:**
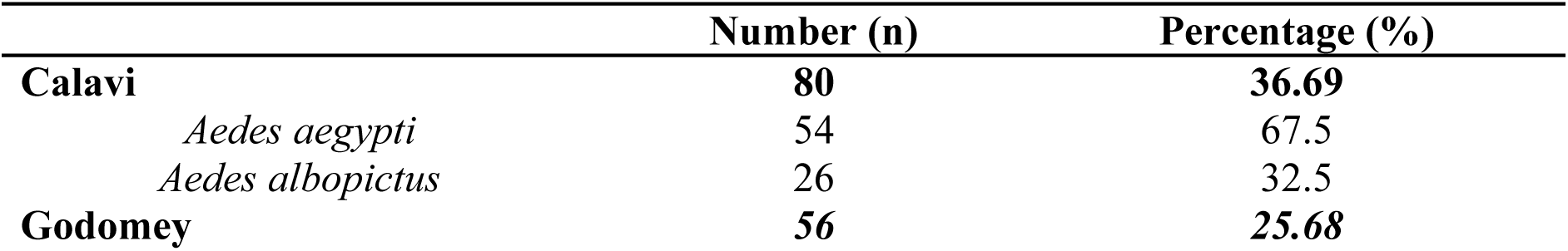
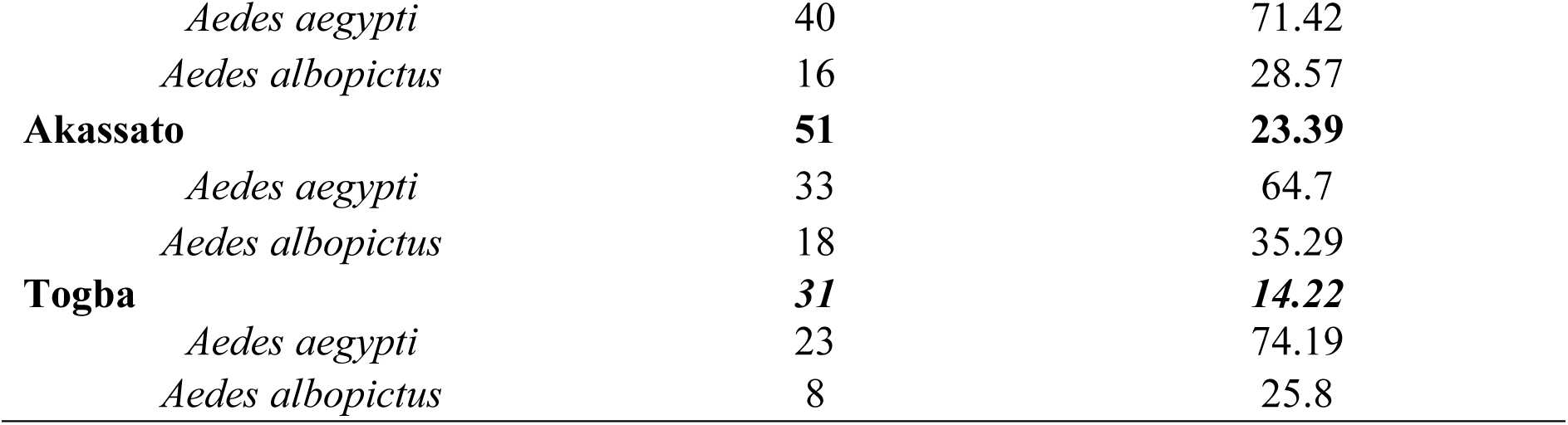
Distribution of *Aedes aegypti* and *Aedes albopictus* mosquitoes in the arrondissement of the commune of Abomey-Calavi.

### 3.2. Prevalence of Dengue virus serotypes in *Aedes aegypti* and *Aedes albopictus* mosquitoes

A total of 218 mosquitoes, including 150 *Aedes aegypti* and 68 *Aedes albopictus*, were analyzed. Analyses revealed the presence of the dengue virus in all arrondissement studied (table II). The analyses show that DENV3 is the most prevalent serotype in *Aedes aegypti*. This serotype is mainly observed in Togba (17.4%) and Godomey (12.5%). Serotypes DENV1 and DENV2 are also present in *Aedes aegypti*, with respective prevalences of 6% and 6.7%. In Togba, DENV2 is more frequent than in the other arrondissement, reaching 13%. DENV4 is the least common serotype and was only detected in *Aedes aegypti* in Akassato (3%) and Togba (4.3%), confirming its low circulation in the mosquito population studied.

**Table II:** Prevalence of Dengue virus serotypes in *Aedes aegypti* and *Aedes albopictus* mosquitoes.

| Study Site | Species | Tested | DENV1 | DENV2 | DENV3 | DENV4 |
| --- | --- | --- | --- | --- | --- | --- |
| Calavi | *Ae.aegypti* | 54 | 4 (7.40%) | 3(5.55%) | 6(11.11%) | 0 |
|  | *Ae.albopictus* | 26 | 0 | 1(3.84%) | 7(26.92%) | 0 |
| Akassato | *Ae.aegypti* | 33 | 2(6.06%) | 2(6.06%) | 3(9.09%) | 1(3.03) |
|  | *Ae.albopictus* | 18 | 0 | 1(5.55%) | 3(16.66%) | 0 |
| Godomey | *Ae.aegypti* | 40 | 3(7.5%) | 2(5%) | 5(12.5%) | 0 |
|  | *Ae.albopictus* | 16 | 2(12.5%) | 2(12.5%) | 0 | 0 |
| Togba | *Ae.aegypti* | 23 | 0 | 3(13.04%) | 4(17.39%) | 1(4.35%) |
|  | *Ae.albopictus* | 8 | 1(12.5%) | 1(12.5%) | 0 | 0 |

Legend: Ae. stands for *Aedes*, and DENV stands for dengue virus.

In *Aedes albopictus* mosquitoes, DENV3 is also the dominant serotype. It is mainly found in Calavi (26.92%) and Akassato (16.66%). DENV1 and DENV2 were detected in Godomey and Togba, with identical prevalences (12.5%). However, no cases of DENV3 were observed in these two arrondissement. Finally, no *Aedes albopictus* mosquitoes carried the DENV4 serotype, confirming its absence in this species within the study area.

### 3.3 Prevalence of Dengue virus serotypes in the studied sites

The analysis of the distribution of dengue virus serotypes in the studied arrondissement revealed variability in infection rates depending on the area (table III). Serotype DENV3 was the most frequent, with an overall prevalence of 12.4% (95% CI: 8.3–17.5%). It was particularly dominant in Calavi (16.2%), Akassato (11.8%), and Togba (12.9%).Serotype DENV2 ranked second with a prevalence of 6.9% (95% CI: 3.9–11.1%), while DENV1 was detected at lower rates (5%, 95% CI: 2.5–8.8%). Serotype DENV4 was the least represented, detected only in the arrondissement of Akassato (2%) and Togba (3.2%), with an overall prevalence of 0.09% (95% CI: 0.1–3.30%).Statistical analysis showed no significant difference in serotype distribution among the arrondissement (p = 0.80).

**Table III:** Distribution of Dengue Virus Serotypes in the Studied arrondissement.

| Arrondissement | Tested | DENV1<br>(%,IC) | DENV2 (%,IC) | DENV3 (%,IC) | DENV4 (%,IC) | P<br>value |
| --- | --- | --- | --- | --- | --- | --- |
| Calavi | 80 | 5 (1.4-12.3) | 5 (1.4-12.3) | 16.2 (8.9-26.2) | 0 |  |
| Godomey | 56 | 8.9 (3-19.6) | 7.10 (2-17.3) | 8.9 (3-19.6) | 0 |  |
| Akassato | 51 | 3.9 (0.5-13.5) | 5.9 (1.2-16.2) | 11.8 (4.4-23.9) | 2 (0-10.4) |  |
| Togba | 31 | 3.2 (0.1-16.7) | 12.9 (3.6-29.8) | 12.9 (3.6-29.8) | 3.2 (0.1-16.7) |  |
| Entire area | 218 | 5 (2.5-8.8) | 6.9 (3.9-11.10) | 12.4 (8.3-17.5) | 0.09 (0.1-3.30) | 0.80 |

Legend: DENV stands for dengue virus.

### 3.4. Evaluation of Dengue Epidemic Risk

The table below presents the assessment of epidemic risk using the House Index (HI) across the arrondissement (table IV). The House Index varies between arrondissement but remains generally moderate. The arrondissement of Calavi recorded the highest index at 18.4% (95% CI: 10.9–28.1). Godomey follows with 16.1% (95% CI: 8–27.7), while Akassato shows the lowest index at 12.5% (95% CI: 5.6–23.2). The overall index for the entire area is 15.5% (95% CI: 11.3–20.6). The risk level remains moderate in all arrondissement, indicating a potential risk of a dengue outbreak.

**Table IV:** Dengue Epidemic Risk Assessment Using the House Index (HI) by arrondissement.

| Arrondissement | I. House (95%IC) | Risk Assessment |
| --- | --- | --- |
| Calavi | 19.04 (10.9-28.10) | moderate |
| Godomey | 16.1 (8-27.7) | moderate |
| Akassato | 12.5 (5.6-23.2) | moderate |
| Togba | 11.90 (4.4-28.1) | moderate |
| **Entire area** | **15.5 (11.3-20.6)** | moderate |

Legend: I. House for House Index; DENV stands for dengue virus; CI stands for confidence interval.

### 3.5. Frequencies of Alleles and Genotypes of *Kdr* Mutations *F1534C, V1016G, S989P, V1016I,* and *V410*L

Five knockdown resistance mutations were detected in *Aedes aegypti* and *Aedes albopictus* collected from four study sites in Benin (table V). The resistant (R) allele frequency for the S989P mutation ranged from 0.333 to 0.509 in *Ae. aegypti* and from 0.333 to 0.500 in *Ae. albopictus*, with significant deviations from Hardy-Weinberg Equilibrium (HWE) observed in Calavi for both species (p = 0.013 and p = 0.006), suggesting possible selective pressure. For the F1534C mutation, R allele frequencies ranged from 0.167 to 0.485 in *Ae. aegypti* and from 0.194 to 0.438 in *Ae. albopictus*, with a significant deviation in *Ae. albopictus* from Togba (p = 0.049). The V1016I mutation showed low R allele frequencies in *Ae. aegypti* (0.125 to 0.287), with a significant deviation in Akassato (p = 0.019), while it was nearly absent in *Ae. albopictus*, particularly in Godomey where all individuals were wild type (SS). The V1016G mutation was more prevalent in Calavi, with R allele frequencies of 0.537 in *Ae. aegypti* and 0.558 in *Ae. albopictus*, both with significant HWE deviations (p = 0.001 and p = 0.004), suggesting ongoing selection. Finally, the V410L mutation had the lowest overall R allele frequency, not exceeding 0.375 in any population, with no significant deviations from HWE except in *Ae. aegypti* from Calavi (p = 0.000), indicating possible strong selection or population structure effects.

**Table V:**
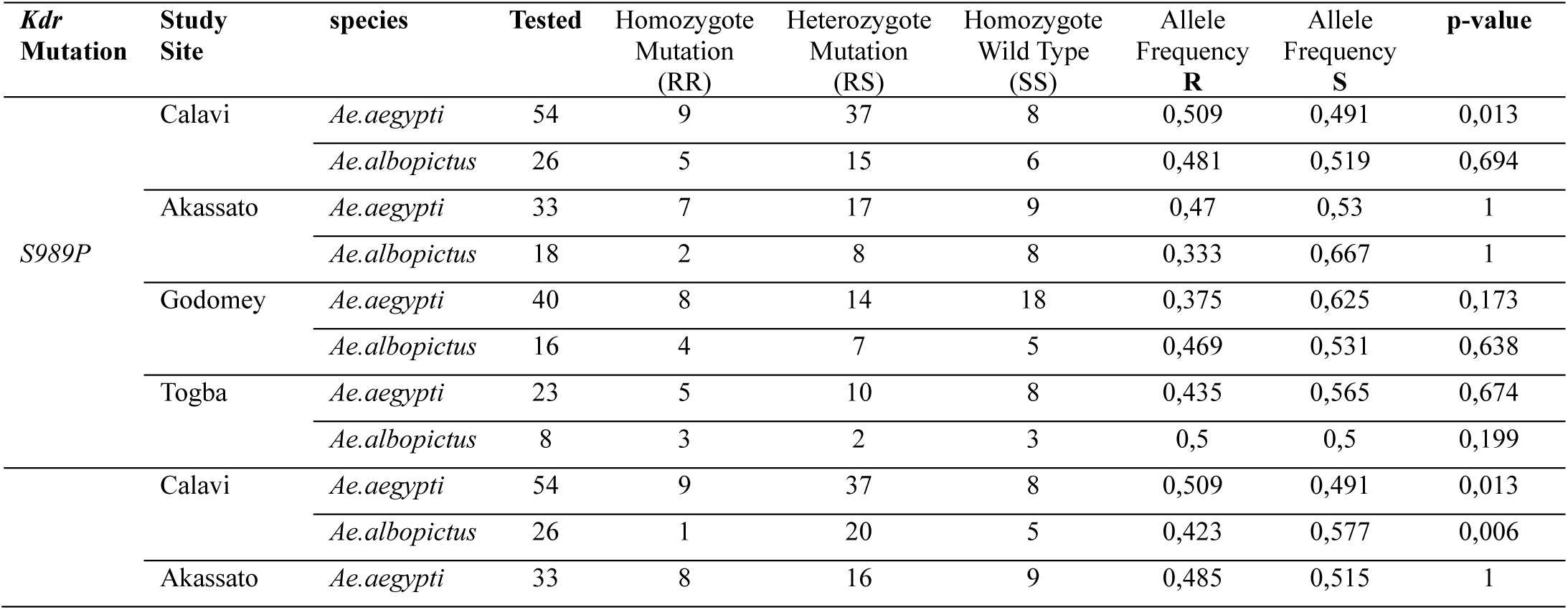
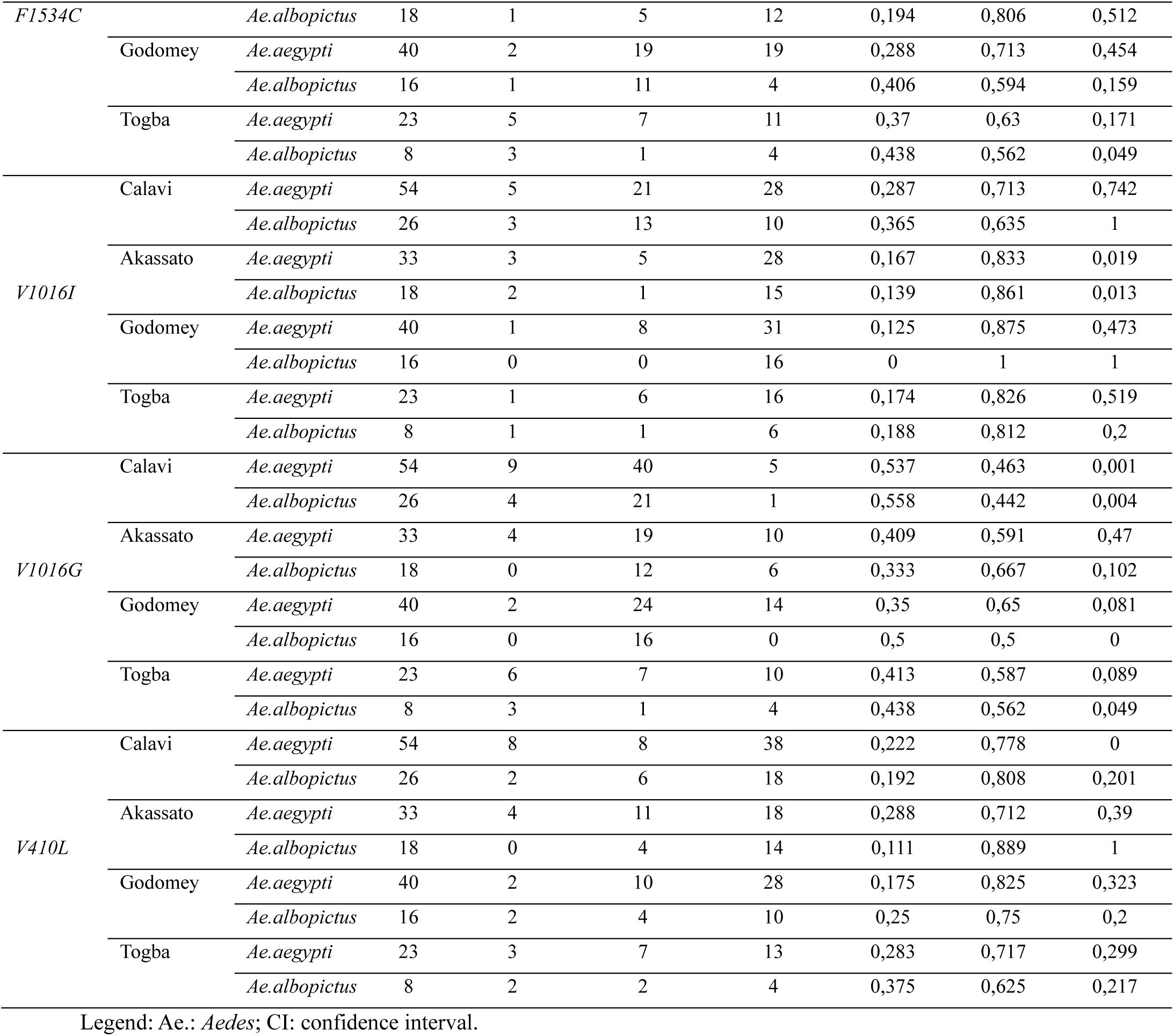
Genotypes and Allelic Frequencies of the Different Mutations.

### 3.6 Distribution of *Kdr* Mutations in Mosquitoes According to RR and RS Genotypes

The analysis of *Kdr* mutation distribution according to homozygous resistant (RR) and heterozygous (RS) genotypes highlights the presence of several mutation combinations. Among RR genotype mosquitoes, 22 individuals carry a double mutation and 4 exhibit three mutations (figure 2). Of the latter, two *Aedes aegypti* individuals carry the *F1534C, S989P*, and *V410L* mutations, while two *Aedes albopictus* mosquitoes respectively carry the *F1534C, V1016G, S989P* mutations and the *F1534C, V1016G, V410L* mutations. As for the RS genotype, 67 mosquitoes carry two mutations, 18 have four, and 3 individuals are carriers of all five studied mutations.

**Figure 2:**
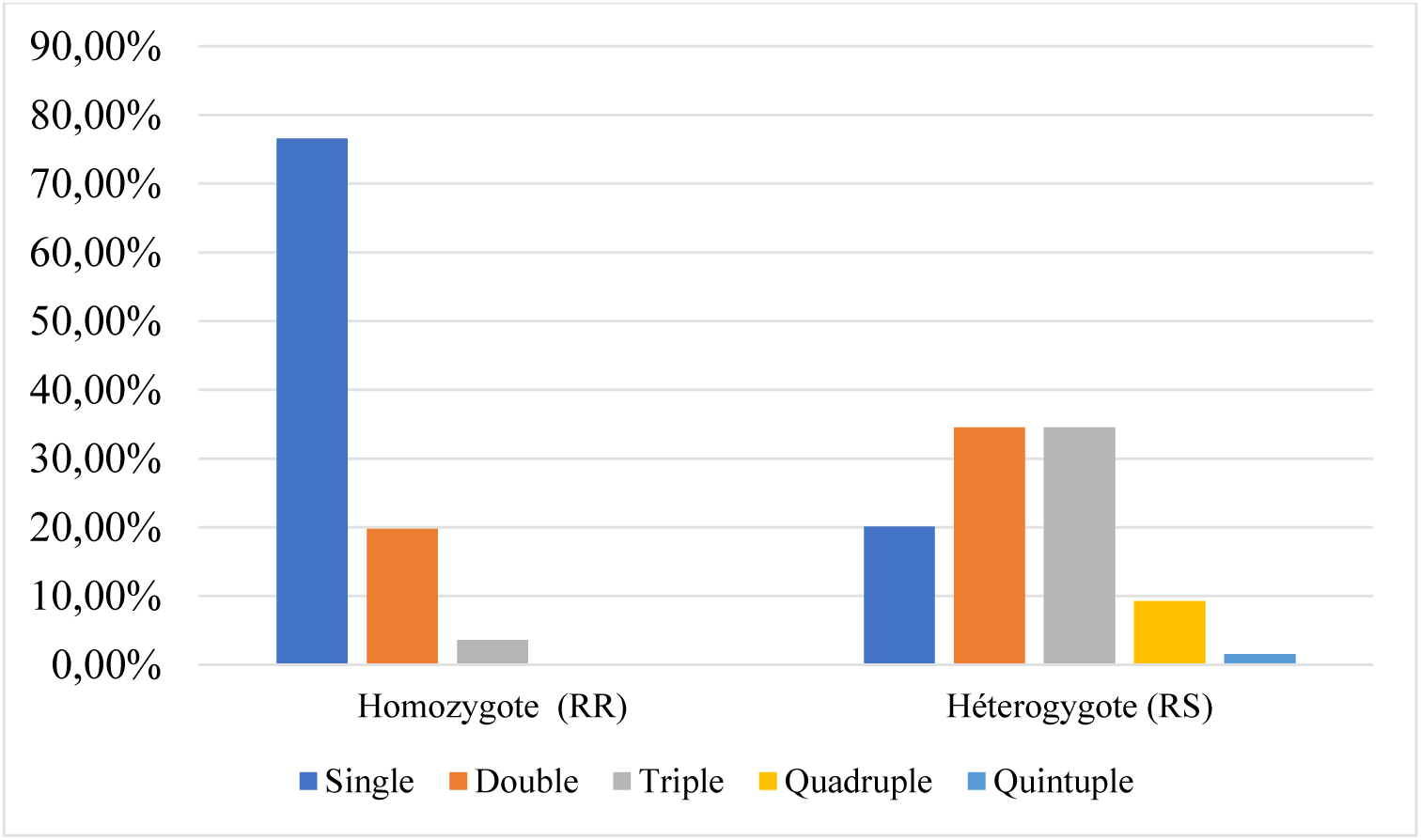
Distribution of Mutations According to Homozygous (RR) and Heterozygous (RS) Genotypes in *Aedes* Mosquitoes in the Study Area

### 3.7 Co-occurrence of Kdr Mutations F1534C, V1016G, S989P, V1016I, and V410L

The study of the co-occurrence of *Kdr* mutations reveals that certain combinations are more frequent than others (figure 3). The most common combination is that of *F1534C* and *V1016G* mutations, found in 11 mosquitoes, suggesting a strong association between these two mutations. The *F1534C* mutation is also frequently associated with *S989P* (6 mosquitoes) and *V410L* (5 mosquitoes). Similarly, the combination of *S989P* and *V410L* mutations is observed in 4 mosquitoes, although this association is less frequent. On the other hand, the *V1016I* mutation was not found in co-occurrence with either *F1534C* or *V410L*, suggesting either a weak association between these mutations or a lower selection pressure favoring these combinations.

**Figure 3:**
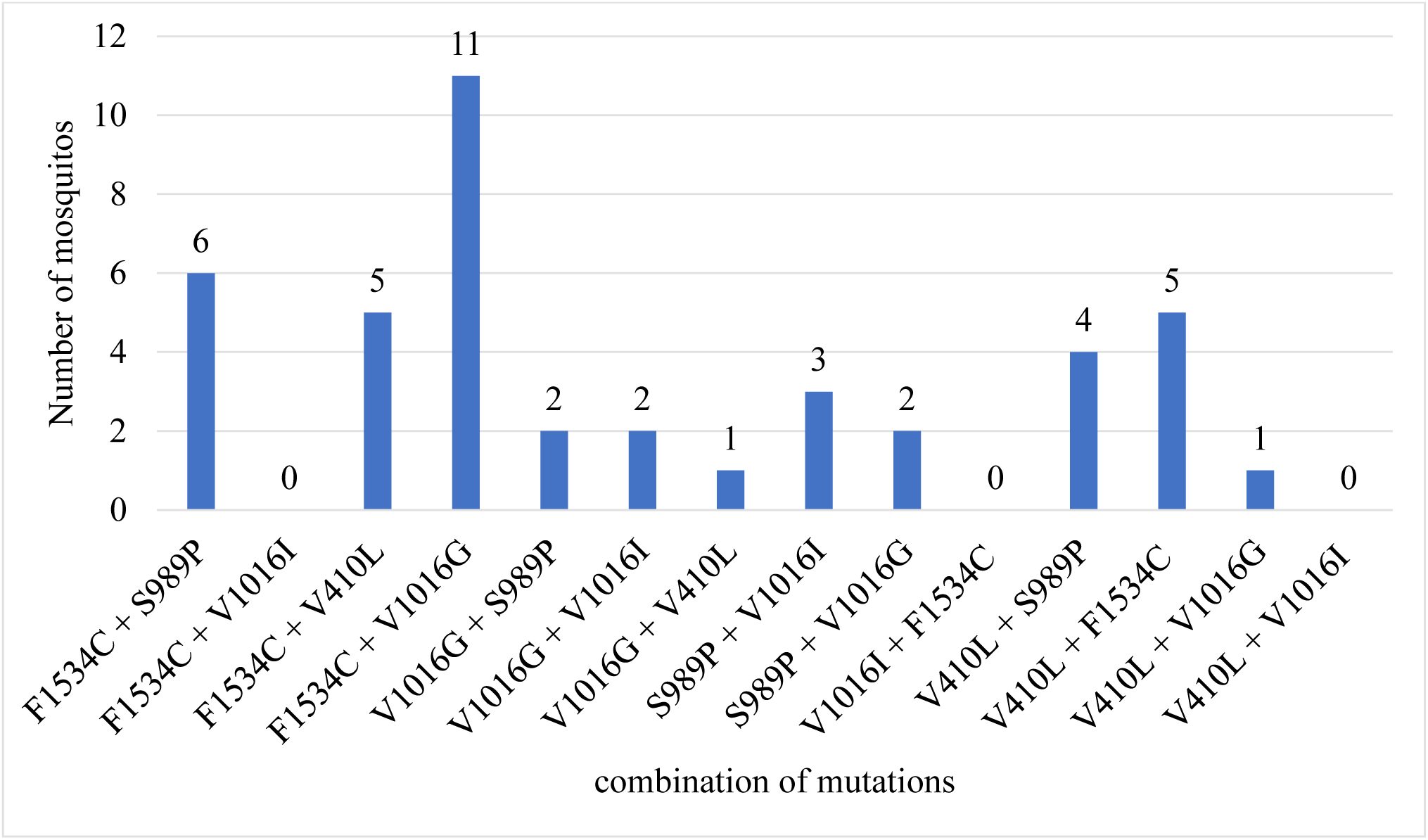
Number of Mosquitoes Based on Two Mutations.

### 3.8. Relationship Between Infection and Carrying of Different *Kdr* Mutations

The analysis of the relationship between dengue infection and the presence of *Kdr* mutations shows the frequency of resistant (RR) genotypes in dengue-positive mosquitoes (table VI). In *Aedes aegypti*, the frequency of homozygous resistant mosquitoes is higher in infected mosquitoes for *F1534C* (17.6% vs 15.5% in negatives), *V1016G* (20.6% vs 12.1%), and *S989P* (26.5% vs 20.7%). In *Aedes albopictus*, the frequencies of homozygous resistant mosquitoes are also higher in dengue-positive mosquitoes: 44% for *F1534C* (vs 33% in negatives) and 53% for *V1016G* (vs 45%). However, the differences observed between positive and negative mosquitoes are not statistically significant (p > 0.05), suggesting that the presence of *Kdr* mutations does not have a direct impact on dengue infection.

**Table VI:** Relationship Between Infection and Carrying of Different Kdr Mutations.

|  |  | *F1534C* | *V1016G* | *S989P* | *V1016I* | *V410L* |
| --- | --- | --- | --- | --- | --- | --- |
| **Positif** |  |  |  |  |  |  |
| *Aedes aegypti* |  |  |  |  |  |  |
|  | RR | 6 | 7 | 9 | 2 | 4 |
|  | RS | 17 | 16 | 15 | 13 | 8 |
|  | SS | 11 | 11 | 10 | 19 | 22 |
|  | Fre (R) | 44 | 44 | 49 | 25 | 24 |
|  | IC_95% | (31-54) | (32-56) | (37-60) | (15-35) | (13-34) |
| *Aedes albopictus* |  |  |  |  |  |  |
|  | RR | 7 | 4 | 3 | 1 | 3 |
|  | RS | 6 | 10 | 8 | 3 | 5 |
|  | SS | 4 | 3 | 6 | 13 | 9 |
|  | Fre (R) | 44 | 53 | 41 | 9 | 32 |
|  | IC_95% | (27-61) | (36-70) | (25-58) | (1-19) | (17-48) |
| **Total** |  |  |  |  |  |  |
|  | RR | 10 | 11 | 12 | 3 | 7 |
|  | RS | 24 | 26 | 23 | 16 | 13 |
|  | SS | 17 | 14 | 16 | 32 | 31 |
|  | Fre (R) | 43 | 47 | 46 | 22 | 26 |
|  | IC_95% | (34-53) | (37-57) | (36-56) | (14-30) | (18-35) |
| **Négatif** |  |  |  |  |  |  |
| *Aedes aegypti* |  |  |  |  |  |  |
|  | RR | 18 | 14 | 24 | 5 | 13 |
|  | RS | 62 | 74 | 51 | 27 | 28 |
|  | SS | 36 | 28 | 41 | 84 | 75 |
|  | Fre (R) | 42 | 44 | 43 | 16 | 23 |
|  | IC_95% | (36-49) | (38-50) | (36-49) | (11-21) | (18-29) |
| *Aedes albopictus* |  |  |  |  |  |  |
|  | RR | 2 | 3 | 11 | 5 | 3 |
|  | RS | 30 | 40 | 24 | 12 | 11 |
|  | SS | 19 | 8 | 16 | 34 | 37 |
|  | Fre (R) | 33 | 45 | 45 | 22 | 17 |
|  | IC_95% | (24-42) | (35-55) | (35-55) | (14-30) | (9-24) |
| **Total** |  |  |  |  |  |  |
|  | RR | 20 | 17 | 35 | 10 | 16 |
|  | RS | 92 | 114 | 75 | 39 | 39 |
|  | SS | 55 | 36 | 57 | 118 | 112 |
|  | Fre (R) | 40 | 44 | 43 | 18 | 21 |
|  | IC_95% | (34-45) | (39-50) | (38-50) | (14-22) | (17-26) |

Legend: CI: confidence interval.

### 3.9. Relationship Between Mosquitoes Carrying Multiple Dengue Serotypes and Multiple Mutations

A total of six mosquitoes were found to carry two serotypes of the dengue virus. No mosquito was found with more than two serotypes. The analysis of these mosquitoes reveals a diversity in the presence of *Kdr* mutations:

- Three mosquitoes were infected with DENV1 and DENV3. Among them, one carried no mutations, while the other two had either the S989P mutation alone or the combination of mutations *V1016I* and *S989P*.
- One mosquito infected with DENV1 and DENV2 carried only the V410L mutation.
- One mosquito carrying DENV2 and DENV3 had no mutations.
- Finally, one mosquito infected with DENV3 and DENV4 carried the mutations F1534C and V1016G.

This distribution shows that no clear pattern emerges between carrying multiple serotypes and the presence of *Kdr* mutations. Some mosquitoes that were multi-infected carried no mutations, while others had one or more mutations.

### 3.10. Relationship Between Dengue Serotypes and Various *Kdr* Mutations

The exploration of the relationship between the dengue virus serotypes and the presence of *Kdr* mutations *F1534C*, *V1016G*, *S989P*, *V1016I*, and *V410L* reveals interesting trends. As shown in Table VII, some mutations seem to be more frequent in certain serotypes. For example, the mutation S989P is more represented in mosquitoes carrying DENV1 (4/12) and DENV3 (7/28), while the mutation *V1016G* is more common in those infected with DENV2 (5/15). In contrast, the mutation *V1016I* is completely absent in mosquitoes infected with DENV2 and DENV4. For DENV4, which was detected in only two mosquitoes, only the F1534C mutation was found. Statistical analyses show that the observed associations are not significant (p > 0.05), suggesting that the distribution of *Kdr* mutations may be independent of the viral serotype. The lack of a statistically significant association between mutations and serotypes further supports the hypothesis that the presence of *Kdr* mutations is not directly influenced by viral infection.

**Table VII:**
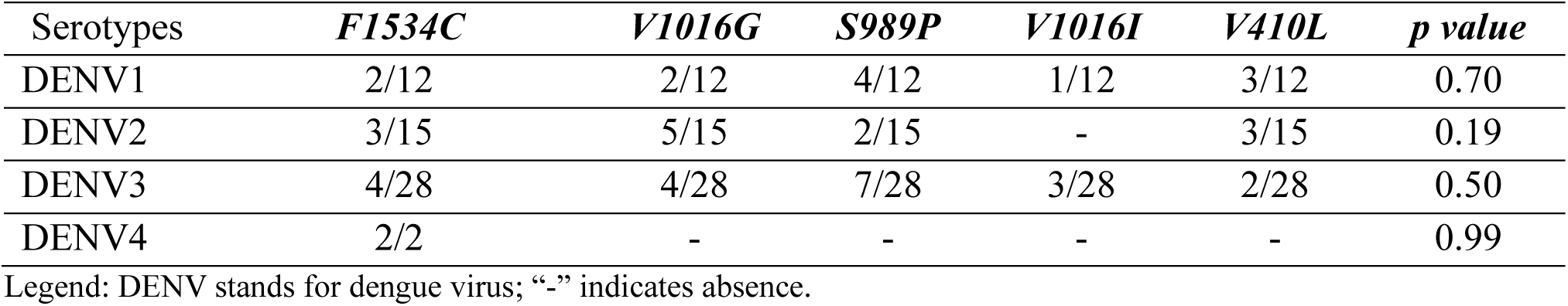
Relationship between the carriage of dengue serotypes and the carriage of different *Kdr* mutations *F1534C, V1016G, S989P, V1016I*, and *V410L*.

| Serotypes | *F1534C* | *V1016G* | *S989P* | *V1016I* | *V410L* | *p value* |
| --- | --- | --- | --- | --- | --- | --- |
| DENV1 | 2/12 | 2/12 | 4/12 | 1/12 | 3/12 | 0.70 |
| DENV2 | 3/15 | 5/15 | 2/15 | - | 3/15 | 0.19 |
| DENV3 | 4/28 | 4/28 | 7/28 | 3/28 | 2/28 | 0.50 |
| DENV4 | 2/2 | - | - | - | - | 0.99 |

Legend: DENV stands for dengue virus; “-” indicates absence.

## Discussion

Longitudinal entomological surveillance of dengue is essential. It helps detect and prevent epidemics while aiding in understanding the spatio-temporal dynamics of the virus. It is also useful for evaluating the effectiveness of vector control strategies. In Benin, the clinical diagnosis of dengue is challenging. It is often confused with malaria due to similar symptoms. Furthermore, there is still no effective early warning system to detect dengue epidemics. The objective of this study was to evaluate the diversity of dengue virus serotypes and the presence of *Kdr* mutations *V410L, V1016G, V1016I, S989P*, and *F1534C* in *Aedes aegypti* and *Aedes albopictus* mosquitoes in the commune of Abomey-Calavi. Our results showed that *Aedes aegypti* mosquitoes, compared to *Aedes albopictus* mosquitoes, were more frequently found in all the studied arrondissement, with a proportion ranging from 64.7% to 74.19%. These observations confirm a previous study conducted in Benin, specifically in 23 communes, where *Aedes aegypti* mosquitoes were found to be the predominant vector, with a proportion of 71.27% [30]. This high presence of Aedes aegypti mosquitoes could be explained by its long history in the region. In contrast, *Aedes albopictus* mosquitoes are more recent introductions [15]. Rapid urbanization and poor waste management contribute to the high presence of Aedes mosquitoes in Benin. The movement of populations and goods, as well as climate change, also play an important role in this proliferation [31]. Several studies have already shown the coexistence of Aedes aegypti and Aedes albopictus mosquitoes in Nigeria, a neighboring country to Benin [32,33].

Our results showed a moderate infection rate in the mosquitoes, with rates varying between 19.6% (CI: 9.8-33.1) and 25% (CI: 16-35.9) depending on the arrondissement. Within the species, the infection rate was 22.7% (CI: 16.2-30.2) in *Aedes aegypti* and 25% (CI: 15.3-37) in *Aedes albopictus*. These results could be explained by the population density in Abomey-Calavi, which favors mosquito proliferation, as well as by Benin’s geographical location. Indeed, the country shares borders with Nigeria and Burkina Faso, both endemic for dengue. Commercial exchanges and population movements between these countries could contribute to the circulation of the virus. No significant difference was observed regarding dengue transmission between *Aedes aegypti* and *Aedes albopictus* mosquitoes. This shows that *Aedes aegypti* mosquitoes are not the only vectors involved in dengue transmission. Several studies have confirmed the involvement of *Aedes albopictus* mosquitoes in dengue epidemics in Africa, suggesting that they should not be neglected in surveillance and vector control strategies [34]. Our results are consistent with a study conducted in six communes of Benin, which revealed the presence of the dengue virus in both *Aedes albopictus* and *Aedes aegypti* mosquitoes captured [30]. However, the overall infection rate for dengue serotypes in our study (23.39%) is much lower than the rate reported by Padonou and colleagues, which was 80%. This difference could be explained by the methodology used. In their study, viral infection was searched in pools of 10 mosquitoes, which would increase the chances of detecting the virus. In contrast, in our study, each mosquito was analyzed individually.

Our results revealed the presence of all four dengue virus serotypes, DENV1, DENV2, DENV3, and DENV4, in the studied arrondissement. Among these serotypes, DENV3 was the most frequent (12.4%, CI: 8.3-17.5), followed by DENV2 (6.9%, CI: 3.9-11.10) and DENV1 (5%, CI: 2.5-8.8). DENV4 was less frequently detected. A similar trend was reported in a previous study evaluating the risk of a dengue epidemic in the Oueme and Plateau departments [35]. However, this study did not detect the presence of DENV2. This serotype, however, was identified in Benin in a study conducted in 2019 on patients suspected of having dengue in the departments of Atlantique, Littoral, and Oueme. This study revealed that 26.7% of the confirmed cases were infected with the DENV2 serotype [11]. Similar results were obtained in Nigeria and Burkina Faso. A systematic study conducted in Burkina Faso to assess epidemiological trends showed that all four serotypes were circulating, with a predominance of DENV3 and DENV2, which aligns with our findings [36]. Additionally, a study conducted in Nigeria in 2019 confirmed the co-circulation of all four serotypes, DENV1, DENV2, DENV3, and DENV4 [37]. These results indicate that Benin is surrounded by countries that are active dengue hotspots, with a high serotypic diversity. Infection by different serotypes can exacerbate the disease and lead to severe forms. This highlights the urgent need to strengthen surveillance and vector control measures in the commune of Abomey-Calavi and throughout the country. Regardless of the arrondissement studied, our results showed a moderate epidemic risk for dengue. This indicates a high risk of dengue serotype transmission in the commune of Abomey-Calavi. Enhanced surveillance is therefore needed to prevent this area from becoming a dengue hotspot in Benin. This situation is all the more concerning in Abomey-Calavi, as deaths due to dengue have already been reported. Similar results have been observed in Benin, with a higher epidemic risk [35].

Currently, prevention is the only effective way to fight against dengue. In our study, we investigated the *Kdr* mutations F1534C, V1016G, S989P, V1016I, and V410L involved in insecticide resistance. Our results revealed that both *Aedes aegypti* and *Aedes albopictus* mosquitoes from Abomey-Calavi carried all of these mutations, with some mutations occurring in co-occurrence, indicating a significant concern regarding the potential impact of insecticide resistance. The causes of mosquito resistance to insecticides are not yet fully understood, but several factors could contribute. The WHO has promoted the use of long-lasting insecticide-treated nets (LLINs) to fight malaria in Africa, a strategy that has yielded good results [38]. In Benin, these nets have been distributed for over a decade with widespread deployment across the country [39]. While this strategy has reduced malaria morbidity and mortality, it has also likely contributed to the development of insecticide resistance in mosquitoes. Frequent use of coils and insecticide aerosols in households may have further reinforced this resistance. However, uncontrolled use of chemical pesticides by farmers has also exacerbated the situation [40]. Agricultural practices involving insecticides to control crop pests exert selective pressure that favors the emergence of resistant populations, not only among pests but also among disease vectors such as those of dengue [41]. Our results showed allele frequencies ranging from 19% (CI: 19-26) to 45% (CI: 40-50). The V1016G mutation had the highest frequency (45%), followed by the F1534C and S989P mutations. These three mutations had already been identified at higher frequencies than our in a study conducted in Godomey and Cotonou by Tokponnon et al., (2024). The frequencies reported in this study for V1016G, F1534C, and S989P were 69%, 64%, and 78%, respectively [23]. The higher resistant allele frequencies and significant deviations from Hardy-Weinberg Equilibrium (HWE) observed in Calavi and Togba for multiple mutations may reflect either strong selective pressure, population structure, or recent gene flow. Similar results were also reported in Nigeria and Ghana. In Ghana, the F1534C mutation was detected at a frequency of 38.73% [42]. In Nigeria, the F1534C and S989P mutations were detected at high frequencies [43]. Our results contrast with those observed in a study in Senegal where the frequencies of mutations V1016I, F1534C, S989P, and V1016G were evaluated. The results of this study showed no frequencies for the mutations sought despite the presence of strong resistance to pyrethroids observed in the *Aedes* mosquitoes included in the study. This difference could be explained by the fact that in Senegal, the resistance observed in the *Aedes* mosquito population is not due to *Kdr* mutations but rather to other mechanisms like metabolic resistance [44]. The V410L mutation was detected in *Aedes* mosquitoes in Angola and Burkina Faso [45,46]. In our study, we identified 22 mosquitoes carrying a double mutation and four carrying three mutations. These results confirm the findings of Tokponnon *et al*., (2024), who showed in their work that 82.9% of pyrethroid-resistant *Aedes aegypti* mosquitoes carried both the *V1016G*, *S989P*, and *F1534C* mutations. This same observation has been made in neighboring countries of Benin. For example, in Nigeria, a study showed that some *Aedes* mosquitoes carried both the S989P and *F1534C* mutations. In Burkina Faso, a *Kdr* mutation association has been observed in *Aedes* mosquitoes since 2019 [46]. The presence of multiple mutations in mosquitoes has also been found in Asia, specifically in China [29]. Our study results revealed a strong association between the *F1534C* and *V1016G* mutations, which were found together in 11 mosquitoes. This co-occurrence of mutations has already been documented in other studies as one of the most common in insecticide-resistant populations. For instance, research conducted in China showed that the co-occurrence of *S989P* and *V1016G* mutations in *Aedes aegypti* is frequently associated with pyrethroid resistance. This association can be explained by their direct impact on sodium channels, which are the primary target of these insecticides [29]. However, in our study, only two mosquitoes simultaneously carried both mutations, suggesting a lower prevalence of this combination in our study area. In Malaysia, the combination of V1016G/F1534C mutations is one of the most widespread, closely followed by the V1016G/S989P combination. It is also interesting to note that in this study, mosquitoes carrying more than four mutations simultaneously were identified, suggesting a progressive evolution of resistance mechanisms due to the selection pressure exerted by repeated insecticide use [47]. These observations highlight the need to diversify vector control strategies, as pyrethroid resistance is well-established in *Aedes* mosquito populations in Benin and several African countries.

One of the main limitations of this study is the inability to conduct sensitivity tests to accurately assess the resistance of mosquitoes to insecticides. While genetic analyses through PCR were performed to detect *Kdr* mutations, these tests do not directly assess mosquito resistance to insecticides under real field conditions. Indeed, phenotypic sensitivity tests, such as contact tests with insecticides, were not conducted, which prevents a more comprehensive evaluation of mosquito populations’ resistance. Although this study provides important information on the serotypic diversity of dengue and mosquito resistance, further research using diversified methods is needed to better understand the extent of resistance and its impact on vector control strategies. Finally, although this study covered several arrondissement in the commune of Abomey-Calavi, the sampling remains relatively limited compared to the geographical diversity of Benin. A larger-scale study, covering more regions of the country, could provide a better estimate of the distribution of dengue and the mutations associated with insecticide resistance on a national level.

## Conclusion

In Benin, dengue transmission is ensured by *Aedes aegypti* and *Aedes albopictus*. This study revealed the mechanism of insecticide resistance through target-site mutations, with the presence of *Kdr* mutations V1016G, V1016I, V410L, F1534C, and S989P at moderate frequencies. These findings suggest a significant level of insecticide resistance, posing a threat to the effectiveness of insecticides and impregnated mosquito nets. The Fisher’s exact test showed no significant difference between the presence of these mutations and the dengue serotypes. Our results highlight the importance of considering the evolution of resistance, which is increasing in frequency and intensity across geographic areas. Therefore, it is imperative to research and develop alternative products and tools to control resistant mosquito populations and limit dengue transmission. Strengthening continuous entomological surveillance and integrating both mosquito species into dengue control strategies is essential. Improved management of larval habitats, along with targeted awareness campaigns, are necessary measures to reduce virus transmission.

## Author Contributions

Conceptualization: T.F.T., O.D.A., Z.S.D., N.K. and M.A.; data collection, T.F.T., O.D.A., Z.S.D., N.K., B.A., O.T., F.G., I.G.Y., F.H.; formal analysis, T.F.T., O.D.A.,Z.S.D., N.K. M.J.A., L.T., R.M.O., R.O and L.M.; mobilization of funding, T.F.T., O.D.A.,Z.S.D., N.K., and R.O.; methodology, T.F.T., R.O., M.A., L.A.M. and M.A.; project administration, T.F.T..; original draft preparation formal, T.F.T., O.D.A.,Z.S.D,, R.M.O, L.A.M. and M.A., supervision, T.F.T., L.B.M, D.K.G and M.A. All authors have read and agreed to the published version of the manuscript.

## Funding

This work was partially supported by researchers from the Entomological Research Center of Cotonou.

Mosquito collections did not directly involve human subjects, but rather their living environments. Permission was requested and obtained before setting up the mosquito traps. All individuals mentioned in this section provided their consent for acknowledgment.

## Acknowledgements

The observations and conclusions presented in this manuscript reflect only the opinion of the author(s). We express our gratitude to the Director of the Centre for Entomological Research of Cotonou, Mr. Germain Gil Padonou, and to his entire team, both in the field and in the laboratory, for their valuable support.

## Data Availability Statement

All data used in this study are included within the article.

## Notes

### Competing Interest Statement

The authors have declared no competing interest.

